# Naïve Huntington’s disease microglia mount a normal response to inflammatory stimuli but display impaired development of innate immune tolerance that can be counteracted by ganglioside GM1

**DOI:** 10.1101/2023.04.05.535712

**Authors:** Noam Steinberg, Danny Galleguillos, Asifa Zaidi, Simonetta Sipione

## Abstract

Chronic activation and dysfunction of microglia have been implicated in the pathogenesis and progression of many neurodegenerative disorders, including Huntington’s Disease (HD). HD is a genetic condition caused by a mutation that affects the folding and function of huntingtin (HTT). Signs of microglia activation have been observed in HD patients even before the onset of symptoms. It is unclear, however, whether pro-inflammatory microglia activation in HD results from cell-autonomous expression of mutant HTT or is the response of microglia to a diseased brain environment, or both. In this study, we used primary microglia isolated from HD knock-in mice (Q140) and wild-type (Q7) mice to investigate their response to inflammatory conditions *in vitro* in the absence of confounding effects arising from brain pathology. We show that naïve Q140 microglia do not undergo spontaneous pro-inflammatory activation and respond to inflammatory triggers, including stimulation of TLR4 and TLR2 and exposure to necrotic cells, with similar kinetics of pro-inflammatory gene expression as wild-type microglia. Upon termination of the inflammatory insult, the transcription of pro-inflammatory cytokines is tapered off in Q140 and wild-type microglia with similar kinetics. However, the ability of Q140 microglia to develop tolerance in response to repeated inflammatory stimulations is partially impaired, potentially contributing to the establishment of chronic neuroinflammation in HD. We further show that ganglioside GM1, a glycosphingolipid with anti-inflammatory effects in wild-type microglia, not only decreases the production of pro-inflammatory cytokines and nitric oxide in activated Q140 microglia, but also dramatically dampen microglia response to re-stimulation with LPS in an experimental model of tolerance. These effects are independent from the expression of interleukin 1 receptor associated kinase 3 (Irak-3), a strong modulator of LPS signaling involved in the development of innate immune tolerance and previously shown to be upregulated by immune cell treatment with gangliosides. Altogether, our data suggest that external triggers are required for HD microglia activation, but a cell-autonomous dysfunction that affects the ability of HD microglia to acquire tolerance might contribute to the establishment of neuroinflammation in HD. Administration of GM1 might be beneficial to attenuate chronic microglia activation and neuroinflammation.

## Introduction

Huntington’s disease (HD) is an autosomal dominantly inherited disorder characterized by neurodegeneration in the corpus striatum, in the cortex and other subcortical brain structures (1). It results from the expansion (> 36) of a stretch of CAG trinucleotide repeats in the first exon of the gene that codes for huntingtin (HTT) (2-4). This mutation translates into an expanded polyglutamine stretch that confers toxic properties to mutant HTT (mHTT) and induces its misfolding and aggregation (5-7).

HTT is ubiquitously expressed throughout the body, but it is particularly abundant in neurons and to a lesser extent in glial cells, including microglia (8-10), where, when mutated, it contributes to various aspects of disease pathology and progression (11-14). Microglia are the myeloid cells of the brain and play a critical role in the normal development and homeostasis of the CNS (15-17). Any alteration of the latter rapidly triggers microglia phenotypic variations according to the specific cues in the extracellular milieu (18-20). Following detection of pathogens or tissue/cell damage, microglia acquire a pro-inflammatory phenotype that involves changes in gene expression and cell morphology, as well as secretion of pro-inflammatory cytokines such as IL-1β, IL-6 and TNF, among others (21). The transient nature of this response is crucial to restoring homeostasis following tissue damage and repair. In many neurodegenerative conditions, however, microglia activation and the production of pro-inflammatory cytokines become chronic and contribute to disease onset and/or progression (22-24).

Pro-inflammatory activation of microglia is already detectable in pre-symptomatic HD patients (25-27), suggesting that it might be an early event in HD pathogenesis, and it correlates with disease progression at later disease stages (28). Whether this pro-inflammatory state is induced by expression of mHTT in microglia in a cell-autonomous manner, by the response of microglia to the ongoing neurodegenerative process (even in prodromic HD), or both remains unclear. Studies in HD animal models have produced conflicting results, as microglia activation and neuroinflammation are present in some but not all models (29-42). Furthermore, most studies have focused on the analysis of microglia morphology and density *in vivo* (31-40), or on the expression of inflammatory cytokines that are not exclusively produced by microglia (12, 31, 36, 38, 43), precluding the possibility to gain direct insights into the specific effects of mHTT expression in microglia.

In this study, we sought to shed light on the cell-autonomous phenotype of HD microglia in the absence of confounding effects from a diseased brain environment. We conducted an extensive analysis of the response of neonatal primary microglia isolated from Hdh140/140 (Q140/140) knock-in mice and wild-type (Q7/7) controls (44) in experimental paradigms that mimic microglia exposure to neurodegenerative conditions *in vivo*, including activation of the toll-like receptors (TLRs) 2 and 4 (45, 46) and exposure to necrotic cells (47, 48). Our studies show that Q140/140 microglia respond to pro-inflammatory stimuli with similar kinetics and strength compared to wild-type microglia. However, Q140/140 microglia fail to develop full tolerance (i.e. to repress pro-inflammatory gene expression (49-51)) in response to repeated LPS stimulations, a dysfunction that could contribute to the establishment of chronic neuroinflammation (52). We further show that treatment of Q140/140 microglia with ganglioside GM1, a glycosphingolipid with anti-inflammatory effects on activated wild-type microglia (53) and with disease-modifying effects in HD mouse models (36), dampens the production of pro-inflammatory cytokines in pre-activated Q140/140 microglia and strengthens tolerance in both wild-type and Q140/140 microglia.

## MATERIALS AND METHODS

### Animal and cells

Primary microglia cultures were prepared from homozygous Q140/Q140 knock-in mice expressing full-length mutant HTT with approximately 140 CAG repeats (54) and from wild-type Q7/Q7 littermates (54). Q140/Q140 mice were kindly donated by Cure HD Initiative (CHDI) and maintained on C57Bl/6J background in our animal facility. All procedures on mice were approved by the Alberta Animal Care and Use Committee and were in accordance with the guidelines of the Canadian Council on Animal Care. Neuroblastoma Neuro-2a (N2a) cells stably transfected with Exon1-mHTT97Q-eGFP or Exon1-wtHTT25Q-eGFP (hereafter referred to as N2a97Q and N2a25Q, respectively) were grown in DMEM (HyClone, SH30022.01): Opti-MEM (Gibco, 31985-070) (1:1) supplemented with 10% fetal bovine serum (FBS, Sigma F1051), 1 mM sodium pyruvate (Gibco, 11360-070) and 2 mM L-glutamine (HyClone, SH30034.01).

### Chemicals and reagents

Ganglioside GM1 purified from porcine brain was provided by TRB Chemedica Inc. (Switzerland) and resuspended in cell culture grade Dulbecco’s phosphate buffered saline (DPBS, HyClone, SH30028.02). Lipopolysaccharide (LPS serotype O55:B5, gamma-irradiated) and lipoteichoic acid (LTA; from *Staphylococcus aureus*) were purchased from Sigma (L6529 and L2515, respectively). Recombinant mouse granulocyte-macrophage colony-stimulating factor (GM-CSF) was purchased from R&D systems (415ML/CF), recombinant human Interferon gamma1b (IFNγ1b) was purchased from Miltenyi Biotec (130-096-481). Exon1-mHTT97Q-eGFP and Exon1-wtHTT25Q-eGFP cloned in pcDNA3.1 plasmids were kindly provided by Dr. Janice Braun (University of Calgary).

### Murine microglia cultures

Primary mixed glial cultures were prepared from P0.5-P1.5 Q140/140 and Q7/7 pups as described previously (55). Briefly, after removal of the meninges, cerebral cortices were enzymatically and mechanically dissociated, and single cell suspensions were seeded in 12-well plates and cultured for 14 days in DMEM/F12 medium (Gibco, 11320-033) supplemented with 10% FBS, 100 U/ml penicillin – 100 µg/ml streptomycin (P/S, Hyclone, SV30010), 1 mM sodium pyruvate (Gibco, 11360-070) and 50 µM β-mercaptoethanol (Sigma, M3148). The cell medium was replaced every 4 days. On day 14-17, cultures were trypsinized as described (55), leaving adherent microglia attached to the bottom of the plate. Microglia were allowed to recover overnight in DMEM/F12 supplemented with 1 mM sodium pyruvate and 50 µM β-mercaptoethanol prior to experiments. Cells were maintained at 37°C in 5% CO_2_.

### Functional studies

After isolation, microglia were cultured overnight in medium without FBS and P/S to let them recover. Microglia were then activated with or without LPS (100 ng/ml) or LTA (10 μg/ml) for the indicated time in fresh serum-free DMEM/F12 medium supplemented with 1 mM sodium pyruvate and 50 μM β-mercaptoethanol. For dose-response experiments with LPS, microglia were incubated with up to 1000 ng/ml LPS for 6 h. In experiments where the decay of the inflammatory response was measured, microglia were first treated with 100 ng/ml LPS for 12 h, then washed once with HBSS^+/+^ (Gibco, 14025) containing 0.1% essential fatty acid-free bovine serum albumin (BSA; Sigma, A8806), once with HBSS^+/+^ and twice more with DMEM/F12, and finally they were cultured in DMEM/F12 medium supplemented with 1 mM sodium pyruvate and 50 µM β-mercaptoethanol for 6-48 h to recover. In experiments that measured the development of tolerance, after stimulation with LPS and recovery in medium without LPS for 24h as described above, cells were reactivated with 100 ng/ml LPS for 6 h. Microglia priming with GM-CSF and IFNγ was performed according to (56). Briefly, one day after isolation, microglia were incubated in DMEM/F12 supplemented with 5% FBS, 1 mM sodium pyruvate, 50 µM β-mercaptoethanol and 5 ng/ml GM-CSF for 4-5 days, with a medium change after the first 48 h. Microglia were then primed for 1 h with 20 ng/ml IFNγ and activated with 100 ng/ml LPS for an additional 48 h. In all experiments where GM1 was used, after stimulation with the indicated factors, cells were washed once with HBSS^+/+^ containing 0.1% essential fatty acid-free BSA, once with HBSS^+/+^ and twice more with DMEM/F12 prior to incubation with 50 μM GM1 in DMEM/F12 supplemented with 1 mM sodium pyruvate and 50 μM β-mercaptoethanol. At the end of each experiment, the conditioned medium was collected for cytokine analysis and centrifuged for 2 min at 800 x g at 4^°^C to remove cellular debris. Cells were lysed in RLT plus buffer (QIAGEN) supplemented with β-mercaptoethanol (10 ul/ml) for RNA extraction, or in ice-cold 20 mM Tris pH 7.4, 1% IGEPAL^®^ CA-630 (Sigma, I8896), 1mM EDTA, 1mM EGTA, 50 μM MG132 (EMD, 474790), 1x phosphatase inhibitor cocktail (PhosSTOP; Roche, 04906837001) and 1x protease inhibitor cocktail (cOmplete; Roche, 04693159001) for protein analysis. Protein concentration in cell lysates was measured with the bicinchoninic acid assay.

### Cell death assay

To measure cellular death, levels of lactate dehydrogenase (LDH) in the culture medium were quantified using CytoTox 96^®^ cytotoxicity assay (Promega, G1780) according to the manufacturer’s instructions. The absorbance at 490nm was read in a SpectraMax i3x (Molecular Devices) and analyzed by SoftMax Pro 6.5.1 (Molecular Devices). Cytotoxicity was calculated as the percentage of total cellular LDH activity released in the conditioned medium. Cell death was also estimated by cell incubation with propidium iodide and high-content microscopy analysis. Briefly, cells were stained with 4µg/ml Hoechst 33258 (Sigma, 861405) for 1 h and then incubated at room temperature with 2µg/ml propidium iodide (Sigma, P4170) for 15 min prior to imaging with MetaXpress (Molecular Devices). Cell death was calculated from the percentage of Hoechst-positive cells that were also stained with propidium iodide.

### Cytokines and nitrite quantification in conditioned medium

TNFα and IL-6 released in the microglia conditioned medium were quantified using mouse TNFα and IL-6 ELISA kits according to manufacturer’s instructions (Invitrogen, 88-7324 and 88-7064-22, respectively) and a SpectraMax i3x spectrophotometer (Molecular Devices). Nitrite levels were measured by the Griess method using sulfanilamide (Sigma, S9251) and N-(1-Naphthyl)ethylenediamine dihydrochloride (Sigma, N9125), as previously described (57). TNF, IL-6 and nitrite levels were normalized over the total protein content in the corresponding cell lysates.

### RNA extraction and qPCR analysis

Primary microglia were collected in RLT Plus buffer and total RNA was isolated and purified with the RNeasy Plus micro kit (QIAGEN, 74034) according to the manufacturer’s instructions. mRNA was reversed transcribed using oligo dT primers (Invitrogen, 18418-012) and SuperScript II (Invitrogen, 18064-014). qPCR was carried out using PowerUp SYBR Green Master Mix (Applied Biosystems) in a StepOne Plus instrument (Applied Biosystems). Unless otherwise indicated, gene expression was normalized over the geometric mean of three reference genes encoding cyclophilin A, ATP synthase F1 subunit beta and Ribosomal Protein Lateral Stalk Subunit P0 (Normalization Index), according to (58).

### Preparation of necrotic N2a cells

Necrotic cells were prepared from N2a25Q and N2a97Q cells. Cells were washed once with HBSS supplemented with 0.1% BSA and once with HBSS, trypsinized with 0.05% Trypsin/0.53 mM EDTA (Corning, 25051CI), spun down at 1,000 x g and resuspended in DMEM/F12 medium supplemented with 1mM sodium pyruvate and 50 µM β-mercaptoethanol at the concentration of 10^7^ cells/ml. Five hundred µl of resuspended cells was transferred into a 15 ml tube and necrotic death was induced by applying 5 freezing-thawing cycles as previously described (59). In each cycle, cells were incubated on dry ice for 2 min followed by thawing in a water bath at 37^°^C for 2 min. The percentage of necrotic cells was assessed using trypan blue staining (HyClone, SV30084.01). Cells were exposed to the freeze-thaw cycle until >98% cell death was achieved. Necrotic cells were immediately incubated with microglia for 4 h at a ratio of 2 necrotic cells per microglial cell.

### Flow cytometry assay

To quantify plasma membrane TLR2 and TLR4 expression, microglia were detached from the culture dishes using StemPro Accutase (Gibco, A1110501), washed with cold HBSS and stained with LIVE/DEAD Fixable Far Red dye (Invitrogen, L34974) for 10 min on ice. Cells were then incubated with mouse FcR Blocking Reagent (Miltenyi Biotec, 130092575) for 10 min at 4°C, followed by incubation with 2 ug/ml PE/Cyanine7-conjugated anti-mouse TLR4 antibodies (BioLegend, 145407) or 2 ug/ml PE-conjugated anti-mouse TLR2 antibodies (BioLegend, 148603) for 30 min at 4°C. Cells were then washed with 0.5% BSA and 2 mM EDTA in PBS, and fixed with 4% paraformaldehyde (Electron Microscopy Sciences, 15710) for 10 min. Samples were stored in 2% PFA at 4^°^C prior to analysis using an Attune NxT Flow Cytometer (Invitrogen) in the Flow Cytometry Core Facility of the Faculty of Medicine & Dentistry at the University of Alberta. Data were analyzed using FlowJo software (version 10.7.1).

### Statistical analysis

Two tailed *t*-test and ratio *t*-test analysis, one-way ANOVA or two-way ANOVA with Tukey’s or Sidak’s multiple comparisons test were performed as indicated in the figure legends, using GraphPad Prism 9. Paired estimation plots (60) of the effects of GM1 treatment on TNF secretion upon induction of tolerance were obtained using an online version of EstimationStats (https://www.estimationstats.com/#/). In each figure, N represents the number of independent experiments performed with different microglia cultures.

## RESULTS

### Q7/7 and Q140/140 microglia respond with similar strength of activation to LPS stimulation

To determine whether HD microglia display cell-autonomous activation and/or exaggerated responses to inflammatory stimuli, we measured the expression of pro-inflammatory cytokines in primary microglia isolated from knock-in Q140/140 and Q7/7 mice, in naïve conditions and after stimulation with LPS, a TLR4 ligand. Naïve microglia of both genotypes had similar plasma membrane levels of TLR4 (Supplementary Fig. 1A.I) and did not express detectable levels of mRNA encoding pro-inflammatory cytokines, including IL-1ϕ3, IL-6 and TNF (Fig. 1A-B). Cell exposure to 100 ng/ml LPS induced upregulation of *Il-1b, Il-6* and *Tnf* mRNA expression (effect of time: *p*<0.05), starting at 3h of exposure, peaking at 9h and then slowly decreasing at 24 and 48h of continuous LPS exposure (Fig. 1A). The gradual decrease in the transcription of pro-inflammatory genes after 9h incubation in LPS was likely due to a physiological weakening of LPS-mediated signaling over time (61) and not to cell death, as there was no difference in cell survival between naïve and LPS-stimulated cells up to 24h, and only a modest increase in the number of dead cells at 48h compared to 24h in Q7/7 microglia (effect of time: *p* = 0.008), but not in Q140/140 microglia (effect of time: *p* = 0.21) (Supplementary Fig. 2). The expression of pro-inflammatory cytokines over time and at each time-point was similar for Q7/7 and Q140/140 microglia, suggesting similar kinetics of TLR-4 activation and downstream target gene transcription. Secretion of TNF (Fig. 1B) and nitric oxide (detected as nitrite in the conditioned medium, Fig. 1C) were also overall comparable between genotypes up to 48h of incubation with

**Fig. 1.**
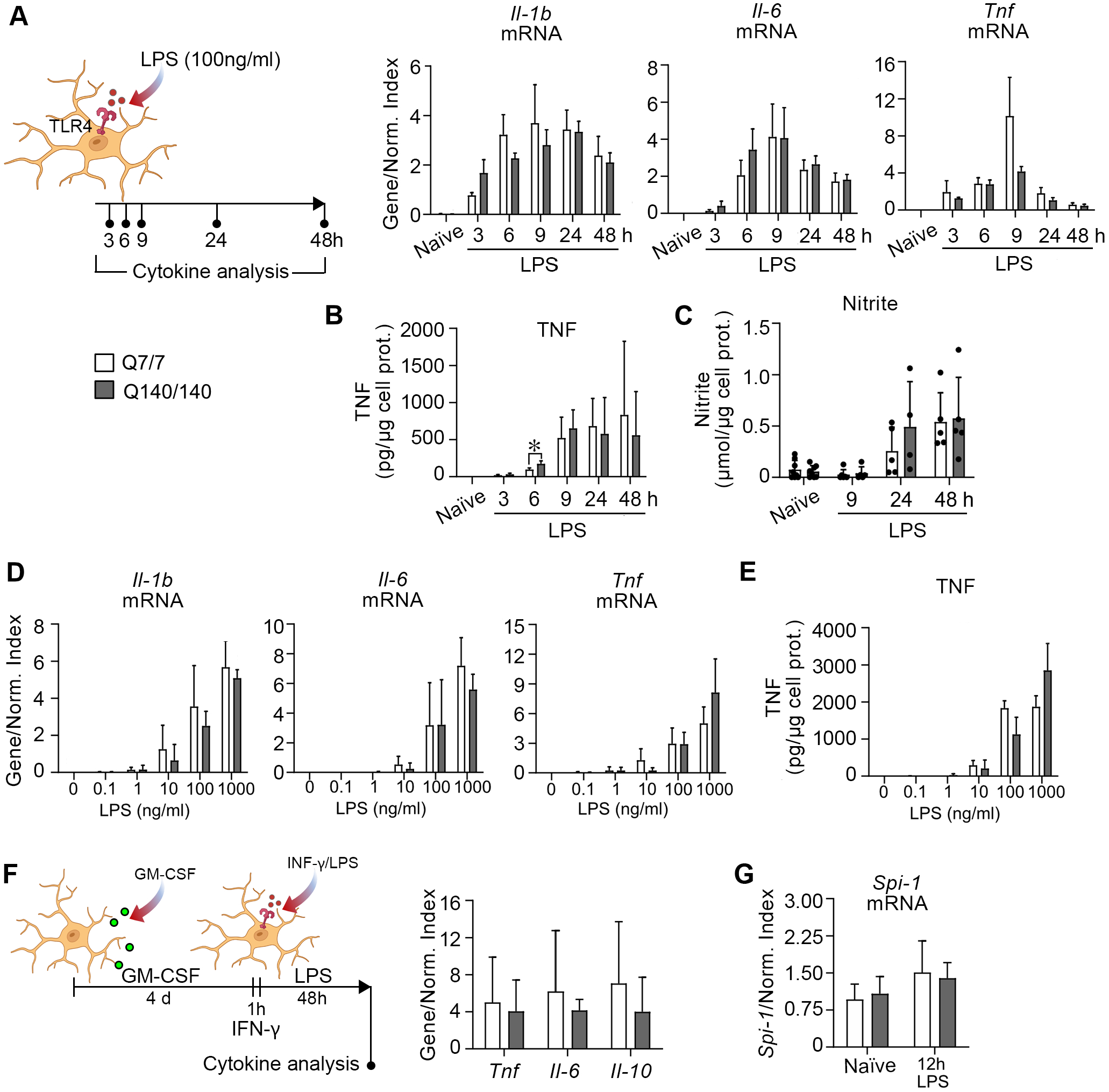
Comparable responses of Q7/7 and Q140/140 microglia stimulated with LPS. **(A)** Microglia were stimulated with LPS (100 ng/ml) for the indicated times. Expression of *Il-1b*, *Il-6* and *Tnf* mRNA at each time-point was normalized over the geometric mean of three housekeeping genes (Normalization Index). N ≥ 3. **(B)** TNF secreted in the conditioned medium was estimated by ELISA. N ≥ 3. **(C)** Nitrite in the conditioned medium. N=8 for naïve microglia, N ≥ 4 for stimulated microglia. No statistically significant differences between genotypes were found for any of the measurements indicated above at any of the time points, except for TFN secretion at 6 h of LPS stimulation. **(D-E)** Dose-response of LPS stimulation. Microglia were stimulated for 6 h with LPS at the indicated concentrations and mRNA was quantified by qRT-PCR. No differences in the expression of *Il-1b*, *Il-6* and *Tnf* mRNA **(D),** and the amount of TNF released in the medium **(E)** were observed between Q7/7 and Q140/140 microglia at all concentrations of LPS tested. N=3 for Q7/7 and N=4 for Q140/140 microglia. **(F)** Microglia were primed with GM-CSF for 4 days, followed by 1 h incubation in IFN-γ and 48 h in LPS to induce a pro-inflammatory state. Expression of *Tnf*, *Il-6* and *Il-10* was normalized as in (A). N ≥ 4. **(G)** mRNA expression of *Spi-1* was measured in control conditions and after stimulation for 12 h with LPS (100 ng/ml) and normalized as in (A). N ≥ 4. In A-F, a one-way ANOVA was used to confirm the effect of treatment in each genotype. The paired two-tailed *t*-test was used to compare differences between genotypes at each time point and LPS concentration. In G, a two-way ANOVA with Tukey’s comparisons test was used. Bars are mean values ± STDEV. *, *p*<0.05; **, *p*<0.01.

LPS. At 6h, the levels of TNF secreted by Q140/140 microglia appeared to be statistically different (and higher) compared to Q7/7 cells. However, considering that at later time points TNF secretion was comparable between genotypes and that mRNA expression was similar at all time points, the biological relevance of this statistical effect is likely negligible.

Next, we sought to determine whether Q140/140 microglia might be more sensitive than Q7/Q7 microglia to a milder stimulation, resulting in a stronger activation at lower doses of LPS. Expression of *Il-1b, Il-6* and *Tnf* (Fig. 1D), and secretion of TNF into the medium (Fig. 1E) did not differ between Q7/7 and Q140/140 microglia at LPS concentrations ranging from 0.1 to 1000 ng/ml (Fig. 1D-E). Finally, we also observed no difference between the pro-inflammatory response of Q140/140 and Q7/7 microglia primed with GM-CSF for 4 days and with IFNγ for 1 h prior to activation with LPS for 48 h, an alternative protocol commonly used to polarize microglia towards a pro-inflammatory state (56) (Fig. 1F).

It was previously suggested that expression of *Spi-1*, a key factor in myeloid fate determination, is upregulated in HD models and mediates cell-autonomous microglia activation and increased microglia response to LPS (12). Consistent with their normal response to LPS stimulation, Q140/140 microglia expressed similar levels of *Spi-1* mRNA as Q7/7 microglia, in both naïve and stimulated conditions (Fig. 1G).

### Q7/7 and Q140/140 microglia show comparable levels of activation following exposure to a TLR2 ligand or necrotic cells

Similar to TLR4, TLR2 is expressed in microglia (62) and contributes to microglia activation in neurodegenerative contexts (62-65). Q7/7 and Q140/140 microglia express similar levels of TLR2 at the plasma membrane (Supplementary Fig. 1B). To investigate the response of Q140/140 microglia to TLR2 stimulation compared to Q7/7 cells, we incubated microglia with lipoteichoic acid (LTA, 10 μg/ml), a TLR2 ligand (61) (Fig. 2A). This treatment resulted in upregulation of *Il-1b*, *Il-6* and *Tnf* transcription (*p*<0.05, Fig. 2B) and secretion of TNF (*p*<0.05) by cells of both genotypes, to a similar extent (Fig. 2C). The levels of nitrite in the conditioned medium after stimulation with LTA were also similar between Q7/7 and Q140/140 microglia cultures (Fig. 2D).

**Fig. 2.**
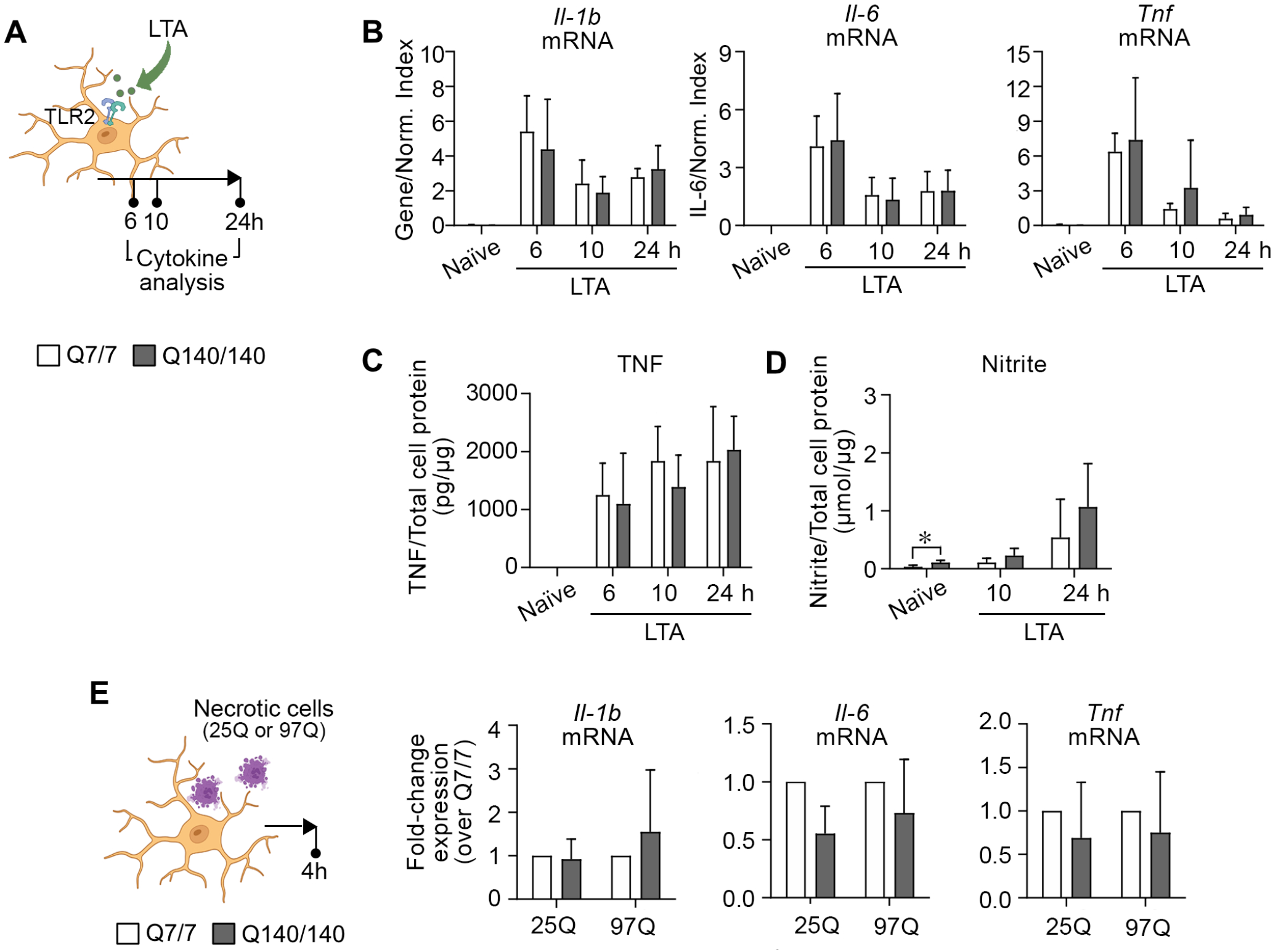
Q7/7 and Q140/140 microglia display similar levels of activation in response to TLR2 ligands and necrotic cells. **(A)** Schematic experimental design showing the time points for cytokine analysis. Q7/7 and Q140/140 microglia stimulated with LTA (10 μg/ml) for 6, 10 and 24 h express similar levels of *Il-1b*, *Il-6* and *Tnf* mRNAs **(B)**, and secrete similar amounts of TNF **(C)** and nitric oxide **(D)** in the medium. Gene expression was normalized over the geometric mean of three housekeeping genes (Normalization Index). One-way ANOVA was used to confirm an effect of treatment in each genotype. Unpaired two-tailed *t*-test was used to compare between genotypes at each time point. N ≥ 3. **(E)** Microglia were incubated with necrotic N2a cells expressing wild-type HTT exon 1(25Q) or mutant HTT exon 1 (97Q) at the ratio of 1:2 (microglia to necrotic cells). mRNA expression of *Il-1b*, *Il-6* and *Tnf* was measured after 4 h by qRT-PCR. Graphs show fold change of mRNA expression in Q140/140 microglia compared to Q7/7 microglia exposed to the same type of necrotic cells. N ≥ 4. Ratio paired *t*-test was used to compare between the genotypes following activation with each specific type of necrotic cell. Bars are mean values ± STDEV. *\*p<0.05*.

To further mimic microglia exposure to activating conditions that are present in neurodegenerative diseases, we incubated microglia with necrotic N2a25Q and N2a97Q cells (Fig. 2E). This resulted in similar levels of activation in Q7/7 and Q140/140 microglia (as measured by the expression of pro-inflammatory cytokine mRNA). Interestingly, incubation of microglia with necrotic N2a97Q cells induced a higher expression of *Il-1b* and *Il-6* compared to exposure to necrotic N2a25Q (Suppl. Fig. 3).

Overall, our data suggest that Q7/7 and Q140/140 microglia respond in a similar manner and with similar kinetics to different types of pro-inflammatory stimuli that are directly relevant to neurodegenerative conditions.

### Q7/7 and Q140/140 microglia display similar kinetics of deactivation after stimulation with LPS

The ability of microglia to cease a pro-inflammatory response when the initial activating stimulus is no longer present is crucial to tissue homeostasis and to prevent chronic microglia activation and tissue damage (52). We exposed microglia to LPS (100 ng/ml) for 12h and then monitored the time required for pro-inflammatory gene expression to return to baseline after the removal of LPS. Within 6h from the end of LPS stimulation, expression of *Il1b, Il-6* and *Tnf* (Fig. 3A) and secretion of TNF (Fig. 3B) returned to baseline in microglia of both genotypes. To confirm this was due to TLR-4 signaling waning off, rather than microglia cell death, we measure the latter at the end of LPS stimulation and 6 and 24h post-stimulation. Microglia cell survival was not affected by the 12h of LPS treatment, although slightly higher levels of lactic dehydrogenase (LDH) in the medium (a surrogate measure of cell death) were detected for Q140/140 microglia in both naïve and stimulated conditions compared to wild-type cells (effect of genotype: F(1,8) = 5.82, *p*=0.04) (Suppl. Fig. 4, 12h LPS). Cell death was minimal (below 10%) and similar in cells of both genotypes even at 24h post-stimulation (effect of treatment: (F1,8) = 5.68, *p*=0.04) (Suppl. Fig. 4), demonstrating that the waning of pro-inflammatory cytokine expression was not due to microglia cell death. Altogether, our data suggest that Q140/Q140 microglia have normal kinetics of deactivation upon cessation of the inflammatory stimulus.

**Fig. 3.**
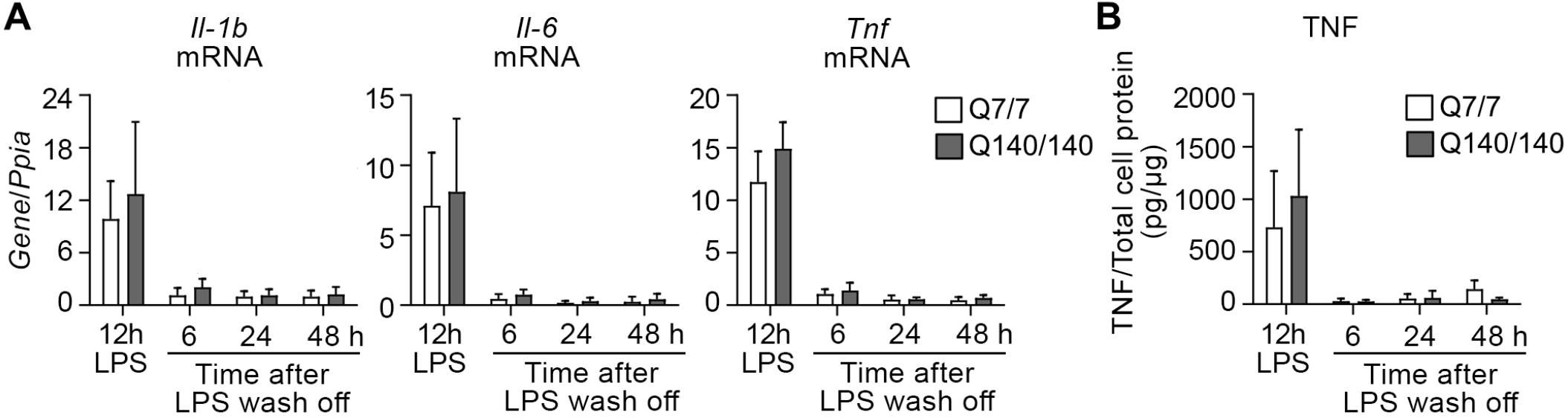
Q7/7 and Q140/140 microglia display similar kinetics of recovery after LPS stimulation. Q7/7 and Q140/140 microglia were pre-incubated with LPS (100 ng/ml) for 12 h, washed and further incubated in serum-free medium for 6, 24 and 48 h. Expression of *Il-1b*, *Il-6* and *Tnf* **(A**, N ≥ 3) and levels of TNF secreted into the medium **(B,** N ≥ 4) were measured at the end of the incubation with LPS (12 h LPS) and during the recovery phase without LPS. mRNA levels were normalized over the housekeeping gene *Ppia* (encoding cyclophilin A). One-way ANOVA was used to confirm the effect of time on microglia phenotype of each genotype. An unpaired two-tail *t*-test was used to compare genotypes at each time-point. Bars are mean values ± STDEV. \**p*<0.05, \*\**p*<0.01, \*\*\*\**p<0.0001*.

### Q140/140 microglia cannot develop full tolerance upon repeated LPS stimulation

Innate immune tolerance is a protective mechanism induced by a preconditioning stimulus that attenuates the inflammatory response to a second stimulation of similar nature (66), including LPS (49-51). It is achieved by epigenetic modifications that silence the expression of pro-inflammatory cytokines and other potentially harmful genes following a first stimulation (67). Similar to peripheral immune cells, microglia can acquire a tolerant state to prevent chronic activation and tissue damage (52, 68). To investigate the ability of HD microglia to develop tolerance, microglia were first stimulated with LPS for 12 h, allowed to recover for 24 h, and then stimulated again with LPS for 6h. Control cells were stimulated only once for 6h (Fig. 4A). As expected, Q7/7 microglia developed tolerance and displayed an attenuated response to the second stimulation. This is clearly shown by the fold-change of pro-inflammatory cytokine expression (Fig. 4B) and secretion (Fig. 4C) relative to cells stimulated only once (dotted horizontal lines in Fig. 4B). In contrast, although *Tnf* expression and secretion (Figs. 4B-C) were attenuated in Q140/140 microglia after repeated LPS stimulation, *Il-1b* and *Il-6* expression were not significantly changed compared to the first stimulation (Fig. 4B), suggesting that these genes were not tolerized in Q140/140 microglia. The mean levels of IL-6 secreted in the medium by Q140/140 microglia after the second LPS stimulation tended to be higher compared to Q7/7 cells (7.39 versus 3.55 pg/mg cell proteins, respectively, Fig. 4C), although the difference between genotypes did not reach statistical significance. Plasma membrane TLR4 levels (Suppl. Fig. 1A.III) and cell viability (Fig. 4D) were similar between Q7/7 and Q140/140 cells, confirming that impaired repression of *Il-1b* and *Il-6* gene expression in Q140/140 microglia was not due to reduced TLR4 signaling or decreased cell viability.

**Fig. 4.**
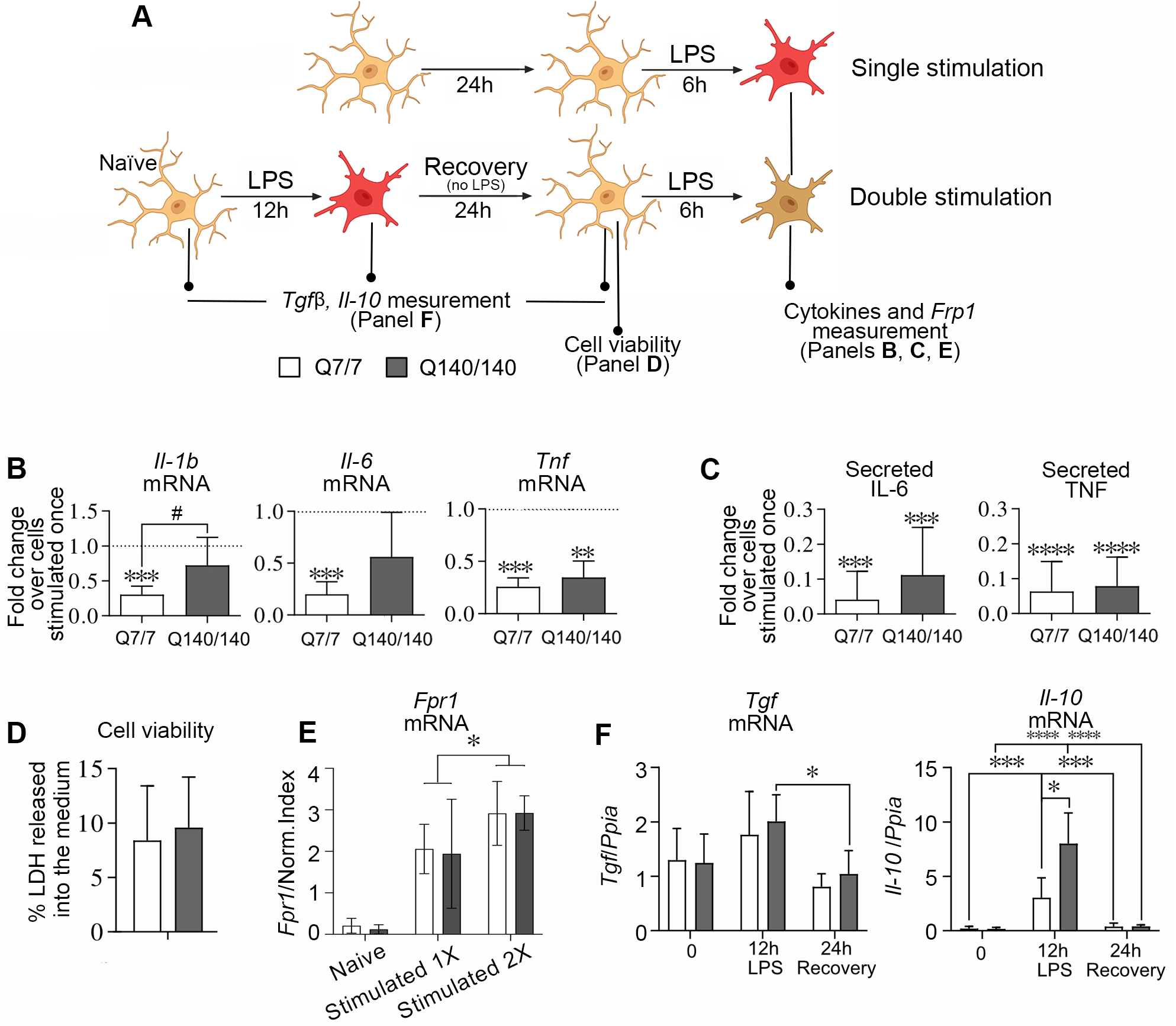
Development of tolerance is partially impaired in Q140/Q140 microglia. **(A)** Schematic experimental design. Microglia were pre-treated with LPS (100 ng/ml) for 12 h, washed and incubated in serum free medium for an additional 24 h (recovery). A second stimulation with LPS (100 ng/ml) was performed for 6 h. Control groups were stimulated only once with LPS for 6h. **(B)** Expression of *Il-1b*, *Il-6* and *Tnf* after the second stimulation with LPS is reported as fold-change compared to cells stimulated only once (baseline represented by the horizontal dotted line in each graph). mRNA levels were normalized over the geometric mean of three housekeeping genes. Ratio *t*-test. Asterisks show significant differences compared to the baseline of cells stimulated only once. The symbol # show a statistically significant difference between genotypes. N 2 5. **(C)** Levels of IL-6 (N 2 4) and TNF (N 2 5) released by microglia in the culture medium. **(D)** Quantification of LDH released in the medium during 24 h recovery period following the first LPS stimulation. Similar levels of LDH released in the medium by Q7/7 and Q140/140 cells indicate microglia were healthy and had similar viability at the time of the second stimulation with LPS. Two-tailed paired *t*-test. N = 3. **(E)** Expression of the non-tolerizeable gene *Fpr1* is increased to a similar extent in Q7/7 and Q140/140 microglia stimulated for a second time with LPS, indicating gene priming. Two-way ANOVA and Sidak’s multiple comparisons post-test. **(F)** *Il-10* and *Tgfb* mRNA expression in naïve microglia, at the end of the first stimulation with LPS and immediately prior exposure to the second dose of LPS (24h recovery). mRNA levels were normalized over cyclophilin A levels. N≥ 4. One-way ANOVA and Tukey’s multiple comparison post-test were used to compare gene expression changes across time points for each genotype. Comparisons between genotypes at each time point were performed with the unpaired two-tail *t*-test. Bars are mean values ± STDEV. *\*p*<0.05*, **p<0.01, ***p*<0.001*, ****p*<0.0001.

While pro-inflammatory genes are tolerized (i.e. repressed) in innate immune cells exposed to repeated stimulations with LPS, other TLR-induced genes involved in anti-microbial activities and tissue repair are not, and might even be primed for faster and increased expression to maintain optimal host defence and tissue homeostasis (67). A prototypical gene that is primed after LPS stimulation is *Fpr1* (formyl peptide receptor 1) (67). Expression of *Fpr1* was higher after the second stimulation with LPS in both Q7/7 and Q140/140 cells, with no significant differences between genotypes (Fig. 4E). Since priming of *Fpr1* depends on TLR4 activation and signaling as much as the silencing of pro-inflammatory genes, these data further confirm that the mechanism underlying impaired tolerance in Q140/140 microglia is downstream of TLR4 activation and is gene-specific.

IL-10 and TGF1β have been involved in the development of tolerance to LPS (69-71). Therefore, we measured the expression of these cytokines after the first stimulation with LPS and during the cell recovery period to determine whether it could account for impaired tolerance in Q140/140 microglia. *Tgfb* expression was comparable between genotypes and not significantly increased after 12h-stimulation with LPS (Fig. 4F). *Il-10* expression was transiently upregulated by LPS in cells of both genotypes, but to a greater extent in Q140/140 microglia compared to Q7/7 (Fig. 4F). Therefore, the impaired ability of Q140/140 microglia to develop full tolerance does not depend on a decreased expression of the tolerogenic cytokines TGFβ and IL-10.

### GM1 has anti-inflammatory effects on activated HD microglia and potently suppresses the expression of pro-inflammatory cytokines when administered prior to LPS re-stimulation

We previously showed that the administration of ganglioside GM1 exerts anti-inflammatory effects on normal microglia activated with various stimuli (53). To determine whether GM1 would have similar effects on HD microglia and could be used to attenuate neuroinflammation in HD, we incubated Q140/140 microglia pre-activated with LPS with 50 µM GM1 for 6h. GM1 decreased the expression of all pro-inflammatory cytokines by Q140/140 microglia (Fig. 5A), like in Q7/7 cells (Suppl. Fig. 5), and dramatically decreased the levels of nitrite in the conditioned medium of cells of both genotypes (Fig. 5B). Similar results were obtained when GM1 was administered to Q140/140 microglia primed with GM-CSF/INFγ and then stimulated with LPS (Fig. 5C), or to microglia stimulated with LTA (Fig. 5D).

**Fig 5.**
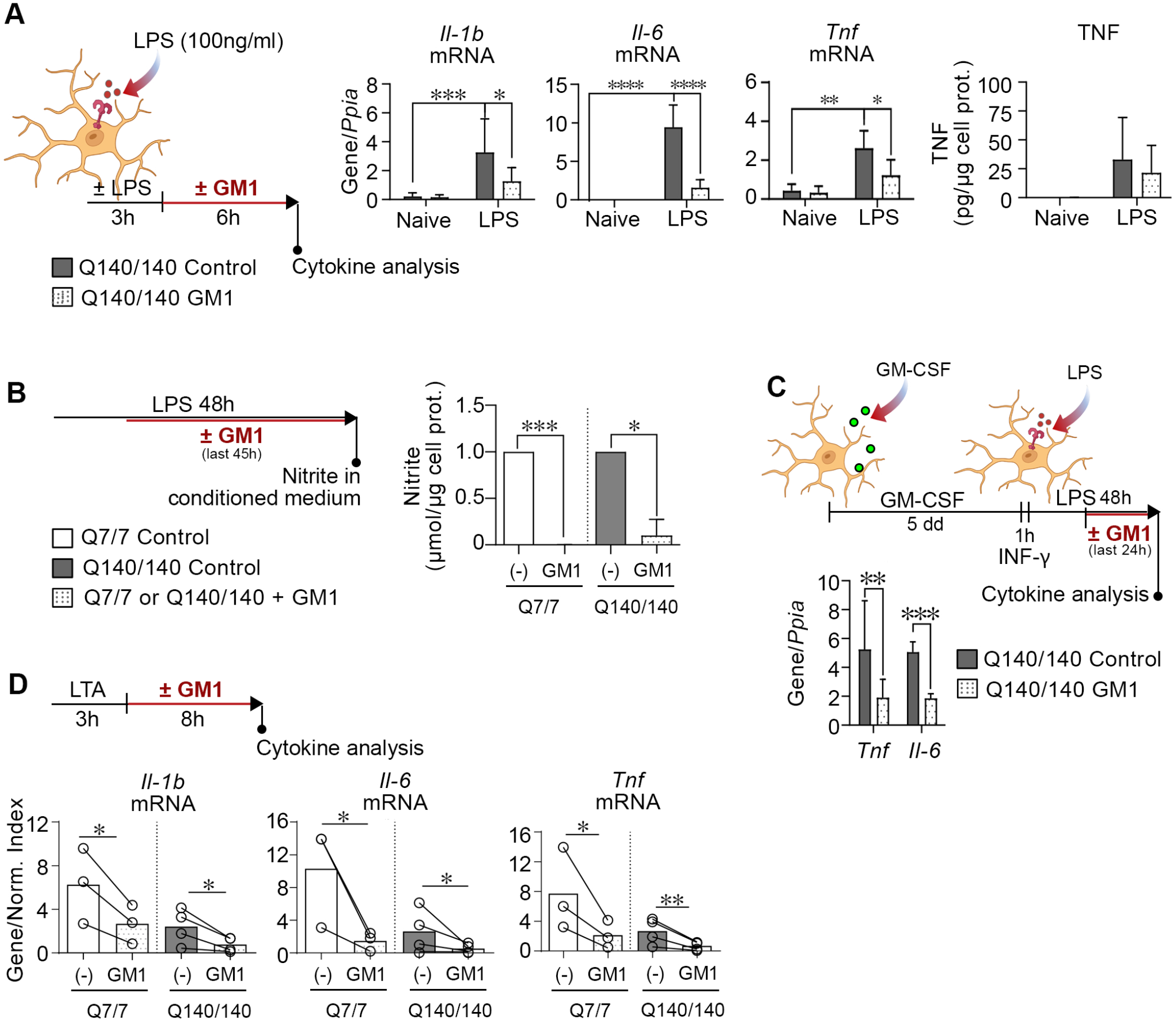
GM1 dampens pro-inflammatory cytokines and NO production in Q140/140 and Q7/7 microglia. **(A)** Schematic experimental design and cytokine measurements. Q140/140 microglia were pre-treated with or without LPS (100 ng/ml) for 3 h, followed by washes and incubation with GM1 (50 µM) or vehicle (PBS) for 6 h. GM1 incubation significantly decreases the expression of *Il-1b, Il-6* and *Tnf* (N 2 4). Gene expression was normalized over the housekeeping gene *Ppia*. TNF secreted in the conditioned medium (N=5) was normalized over the total cell protein content of the cells in each well. Two-way ANOVA with Tukey’s multiple comparisons test. *\*p*<0.05*, **p*<0.01*, ***p*<0.001*, ****p*<0.0001. **(B)** Microglia were treated with 100ng/ml LPS for 3h, followed by incubation with or without GM1 for 45h. Levels of nitrite in the conditioned medium were normalized to total cell proteins. N 2 3. Two-way ANOVA with Sidak multiple comparison test. **(C)** Q140/140 microglia were polarized towards a pro-inflammatory phenotype by incubation with GM-CSF for 4 days, followed by 1h priming with INF-ψ and 48h stimulation with LPS. GM1 or control vehicle were added for the last 24h of incubation in LPS. Gene expression was normalized over *Cyclophilin A*. Ratio paired *t*-test. **(D)** Q7/7 and Q140/140 microglia were incubated with or without LTA (10 ug/ml) for 3 h, followed by washes and treatment with vehicle or GM1 (50 µM) for 8 h. GM1 significantly reduces the expression of *Il-1b*, *Il-6* and *Tnf* mRNA. Gene expression was normalized over the geometric mean of three housekeeping genes (Normalization Index). Ratio paired *t*-test. Bars are mean values ± STDEV. *\*p<0.05, **p<0.01, ***p<0.001*.

Next, we investigated whether treatment with GM1 could restore normal tolerance in Q140/140 microglia. Microglia treatment with the ganglioside during the recovery period (24h) between the first and the second stimulation with LPS (Fig. 6A) resulted in an almost complete abolishment of pro-inflammatory cytokine expression and TNF secretion in both Q7/7 and Q140/140 cells (Figs. 6B-C). Expression of the non-tolerizeable gene *Fpr1* was also dramatically decreased compared to cells that were not pre-incubated with GM1 (Fig. 6D).

**Fig. 6.**
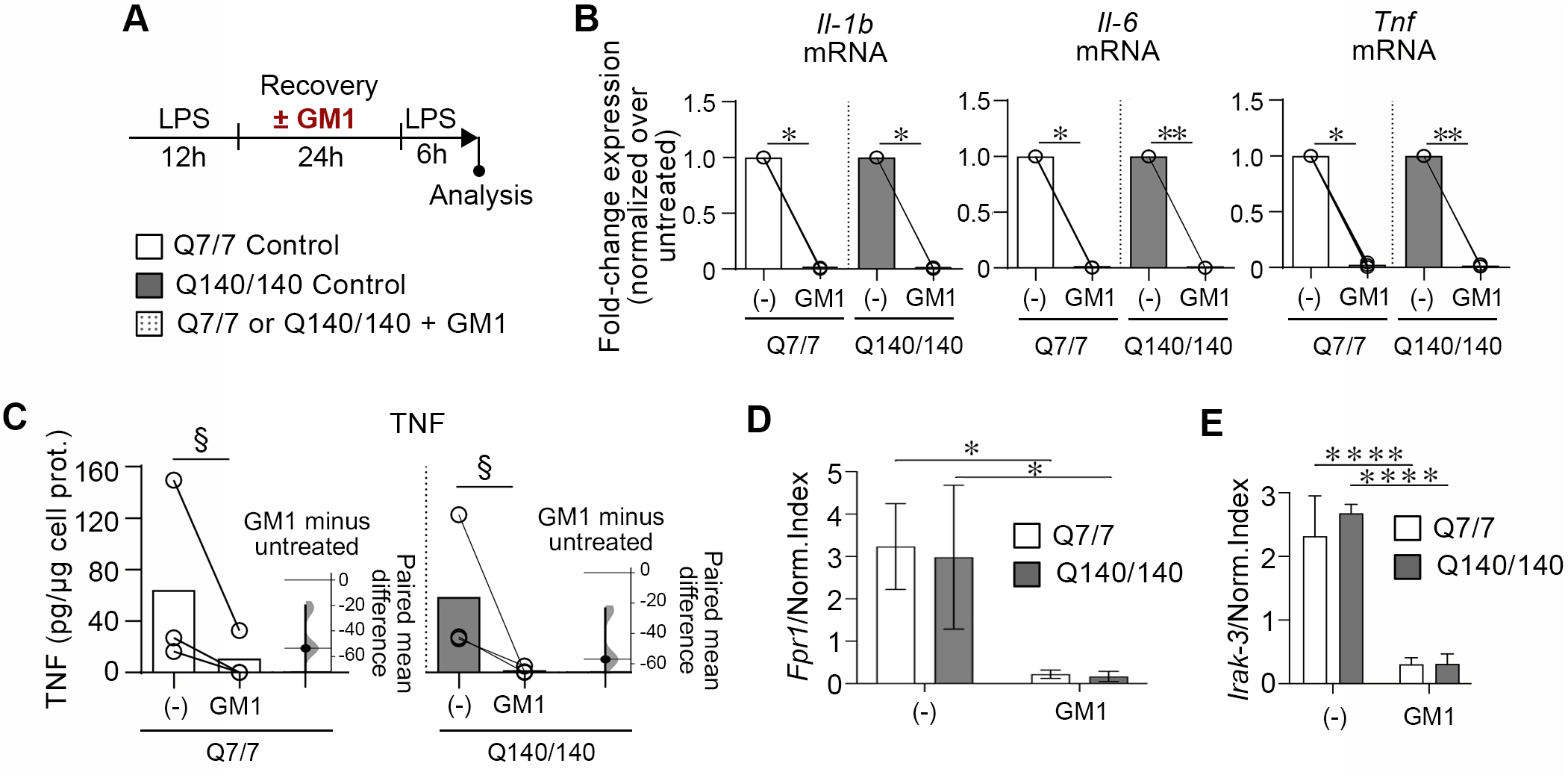
GM1 dampens microglia reactivation in an experimental model of tolerance. **(A)** Experimental design. Microglia were stimulated with LPS (100 ng/ml) for 12 h, then washed and let recover in serum-free medium for 24 h in the presence or absence of GM1 (50 µM). After several washes to remove GM1, a second stimulation with LPS (100 ng/ml) was performed for 6 h, at the end of which, cytokine expression and secretion were measured. **(B)** Expression of *Il-1b, Il-6* and *Tnf* mRNA was normalized over the geometric mean of three housekeeping genes (Normalization Index). N = 3. Ratio paired *t*-test. **(C)** TNF secreted in medium. Data are presented as a paired estimation plot including individual data points (independent experiments), mean values shown by bars, and the bootstrap 95% confidence interval for the effect size. N = 3. §, *p*=0 as per paired estimation plot. **(D)** *Fpr1* expression was measured in microglia stimulated twice with LPS as per experimental design in (A). GM1 presence during the recovery period represses the expression of the non-tolerizeable gene *Fpr1*. **(E)** Cells were treated as in (A). The expression of Irak-3 is similar in Q7/7 and Q140/140 cells. The presence of GM1 during the recovery period prevents upregulation of Irak-3 after re-stimulation with LPS. Two-way ANOVA with Sidak’s multiple comparison post-test was conducted in D and E. Bars are mean values ± STDEV. *\*p<0.05, **p<0.01,* \*\*\*\**p<0.0001*.

Incubation of peripheral monocytes with exogenous gangliosides was shown to cause upregulation of the interleukin-1 receptor associated kinase (IRAK)-M (also known as IRAK-3)(72), an established mediator of innate immune tolerance (73-75). Therefore, we measured the expression of *Irak-3* to determine whether tolerance impairment in Q140/140 microglia and the dramatic effects of GM1 might be mediated by this protein. Expression of *Irak-3* was similar in tolerized Q7/7 and Q140/140 microglia (Fig. 6E) and, contrary to our expectations, it was dramatically decreased when cells were incubated with GM1 prior to a second stimulation with LPS (Figs. 6A and E). Naïve microglia treatment with GM1 did not affect the expression of *Irak-3* (Supplementary Fig. 6). Altogether, our data suggest that microglia pre-treatment with GM1 potently prevents microglia reactivation with LPS and dampens inflammatory responses with a mechanism that is independent of the expression of *Irak-3*.

## DISCUSSION

In this study, we sought to determine whether mHTT expression in HD microglia results in aberrant responses to inflammatory stimuli and abnormal microglia activation in a cell-autonomous manner. Previous work in HD mouse models has often produced inconsistent results. Increased microglia density and/or elevation of pro-inflammatory cytokines was shown in some studies (37-42) (12, 38, 42, 76, 77), but not in others (29-36), even when the same animal models were investigated. Inconsistencies across *in vivo* studies might not be surprising considering genetic differences and potential major confounding effects from environmental conditions and cell-cell interactions in the brain. Studies in isolated neonatal microglia from the same mouse models, however, have produced similarly inconsistent results, with some studies showing higher activation of microglia from R6/2 mice compared to wild-type microglia even in naive conditions (12, 76), and others showing higher activation only after microglia priming (43, 77).

In this study, we used microglia isolated from Q140/140 knock-in mice that express full-length mHTT from the endogenous mouse *Htt* gene locus, a model that closely resembles the genetic makeup of the human disease (44). We demonstrated that Q140/140 microglia display kinetics of activation and deactivation in response to a range of pro-inflammatory stimuli that are remarkably like those in wild-type Q7/7 microglia. To mimic conditions that activate microglia in the context of neurodegenerative diseases, we exposed Q140/140 and Q7/7 microglia to ligands of TLR-4 and TLR-2, two pattern recognition receptors that can promote microglial-mediated neuronal injury and neurodegeneration (78, 79) upon stimulation by pathogenic misfolded proteins and/or endogenous danger-associated molecular patterns (DAMPs) released by injured and dying cells (64, 65, 80, 81). In response to TLR-4 and TLR-2 stimulation, Q140/140 microglia upregulated the expression of pro-inflammatory genes such as *Il1-b, Il-6 and Tnf* and produced NO and TNF to a similar extent and similar kinetics as wild-type cells. These results are in contrast with a previous report where R6/2 microglia stimulated with LPS displayed higher activation compared to wild-type microglia (77). Since the same concentration of LPS and a similar timeframe of stimulation were used in those experiments and in ours, the contrasting results might be due to differences in the genetic makeup of the models used (endogenous levels of expression of full-length mutant HTT in our studies *versus* transgenic expression of a more toxic mutant HTT-exon 1 fragment in R6/2 mice). In a different study, R6/2 microglia were shown to respond more strongly to LPS when primed with IFNγ (43). In our experiments, even when primed with IFNγ Q140/140 microglia upregulated pro-inflammatory gene expression to the same extent as wild-type cells. The kinetics of microglia activation - i.e. the response to different doses of LPS and the time course of pro-inflammatory gene transcription and NO production - were similar between the two genotypes. Q140/140 microglia also retained a normal ability to taper off pro-inflammatory gene transcription upon removal of LPS.

To further mimic conditions that are present in a degenerating brain environment, Q140/140 microglia were incubated with N2a necrotic cells that expressed either wild-type or mutant HTT. Although neuronal death in HD and other neurodegenerative diseases predominantly occurs through pathways other than necrosis (48), stimulation with necrotic cells remains an established model to mimic microglia activation in neurodegenerative conditions where inflammatory intracellular components are released by cells (82-84). Once again, Q140/140 and Q7/7 microglia reacted similarly to incubation with necrotic cells. Altogether, our experiments suggest that expression of full-length mutant HTT at physiological levels does not lead to cell-autonomous activation of murine microglia *per se*, nor to an exacerbated response to pro-inflammatory stimuli in naïve microglia.

Previous studies suggested that increased microglial expression of the myeloid lineage-determining factor PU.1 (encoded by the *Spi-1* gene) - a master regulator of microglia development and function - might underly an exacerbated response to pro-inflammatory stimulation in HD microglia from knock-in (Q175) and fragment (R6/2) HD models (12). In line with our finding of a normal response of Q140/140 microglia to pro-inflammatory stimulation, we also found that *Spi-1* expression was similar in Q140/140 and Q7/7 microglia. These data also confirm our previous observation of normal *Spi-1* expression in the brain of R6/2 mice (36). We speculate that previously reported increases in the expression of *Spi-1* in HD microglia might have resulted from non-cell autonomous microglia exposure to environmental triggers, including pathogens and/or gut microbiota that could indirectly cause microglia priming *in vivo* (85). This appears to be even more likely considering the emerging evidence of a gut-immune system-brain axis with the potential to modulate neuroinflammation and behaviour in HD and other neurodegenerative disorders (86-88).

In our experiments, we noticed that incubation with necrotic cells carrying mutant HTT (97Q) induced a higher expression of *Il-1b* and *Il-6* compared to necrotic cells expressing wild-type HTT (25Q). This suggests that along with classic DAMPs and other mediators of inflammation, HD cells might release additional factors that increase microglia activation, including mutant HTT itself. In support of a higher inflammogenic potential of necrotic HD cells, N2a cells carrying mutant HTT were previously shown to produce higher levels of inflammatory molecules such as MCP-1 and IL-6 compared to wild-type N2a cells (33).

Innate immune tolerance is developed as part of the brain response to acute injuries such as ischemia (89-91) and might limit brain damage and chronic inflammation in neuroinflammatory and neurodegenerative conditions (51, 66, 92). In our experiments, Q7/7 microglia were able to acquire a tolerant state characterized by a lower expression of pro-inflammatory cytokine upon a second exposure to LPS. In contrast, in Q140/140, only TNF was silenced, but not IL-1β and IL-6, the expression of which remained as high as after the first stimulation with LPS. The fact that the transcription of *Il-1b* and *Il-6* was affected differently from *Tnf* is not surprising, since chromatin remodelling and epigenetic marks that lead to gene silencing in tolerant cells are gene-specific (67), and the *Il-1b* gene is less sensitive to the establishment of endotoxin tolerance compared to the *Tnf* gene (93). TLR4 expression at the plasma membrane, which could potentially affect the strength of the tolerogenic signaling, was normal in Q140/140 microglia. Furthermore, priming of the *Fpr1* gene, another consequence of multiple LPS stimulations (67), was not affected in Q140/140 cells, confirming that TLR-4 activation occurred as expected. Other potential players in the development of tolerance, such as the tolerogenic cytokines IL-10 and TGFβ (68, 94, 95) were not likely to be involved either, since their expression in Q140/140 microglia was similar or even higher (for IL-10) compared to Q7/7 microglia. Therefore, the reason underlying incomplete or impaired tolerance in Q140/140 microglia remains to be determined. Perhaps, altered epigenetic mechanisms induced by mutant HTT might interfere with the specific epigenetic modifications, including histone deacetylation and H3K4 demethylation, that drive gene silencing and the development of tolerance (67). Expression of mHTT is indeed associated with epigenetic modifications in hundreds of gene, although most of these tend to repress gene expression (96, 97) and, therefore, would not explain why Q140/140 microglia failed to silence *Il-6* and *Il-1b* in our experiments.

Altogether, our data suggest that an impairment in the ability of HD microglia to develop tolerance, rather than cell-autonomous spontaneous activation or stimulus-induced overactivation of naïve HD microglia, might contribute to a chronic inflammatory state in HD. Impaired tolerance might also explain some of the inconsistencies across different studies in the detection of HD microglia activation, as it could confound data interpretation in animal models exposed to pathogens or dysbiosis. On the other hand, in a pathogen-free environment, the inflammatory potential of microglia that express wild-type or mutant HTT might be similar, at least at early disease stages and in the absence of activating DAMPs, a hypothesis that is also supported by studies that showed that selective depletion of mutant HTT in microglia of BACHD mice did not affect mouse phenotype and pathology (98).

Recently, O’Regan et al. (99) evaluated the inflammatory phenotype of microglia-like cells differentiated from isogenic human iPSCs expressing HTT with polyQ expansions of various lengths. They reported that microglia-like cells with a polyQ expansion (81Q) that is usually linked to juvenile HD, expressed higher levels of IL-6 and TNF following LPS stimulation (1 µg/ml) compared to cells bearing HTT with a normal polyQ length (30Q). However, in cells expressing mutant HTT with 45Q (resulting in adult-onset HD), the secretion of these cytokines was not significantly different from control cells (30Q) (99). Therefore, it is possible that HD microglia might have a higher inflammogenic potential in the context of juvenile HD, but not in adult-onset HD, but further studies with a much larger number of iPSC lines would be required to test this hypothesis. Q140 mice and similar models that express full-length mHTT more closely mirror adult-onset HD, in spite of the larger CAG expansions they carry in their *Htt* gene.

Even if not caused by cell-autonomous effects of mutant HTT expression, microglia activation and neuroinflammation increase with disease progression in HD (28), are likely to contribute to pathology (100) and are a potential target for intervention. In support of this hypothesis, a few studies showed that decreasing glia activation and production of pro-inflammatory cytokines has beneficial effects in HD mouse models: the knock-out of TLR2 or TLR4 extended the life-span of N171-82Q mice(45), a model that overexpresses an N-terminal fragment of mutant HTT in neurons only (101), while inhibition of IKK and the NFkB pathway (102) or TNF signaling (77)in R6/2 mice decreased neurodegeneration and improved mouse behaviour.

We previously showed that the production of pro-inflammatory cytokines by normal murine and human microglia is modulated by endogenous gangliosides and can be drastically decreased by administration of ganglioside GM1 (53). Gangliosides are glycosphingolipids present at the plasma membrane of all cells and are particularly abundant in the brain (103). They play many roles as modulators of cell signaling and immune functions (104, 105), and have profound neuroprotective and disease-modifying effects in HD models (36, 106, 107), where levels of GM1 and other gangliosides were found to be decreased (107, 108). Therefore, we set out to determine whether GM1 could also exert its anti-inflammatory activity on HD microglia. Treatment of HD microglia with GM1 significantly decreased the expression of pro-inflammatory cytokines and the production of reactive nitrite following microglia stimulation with LPS and LTA. Of note, GM1 decreased the expression of all major pro-inflammatory cytokines upon microglia exposure to repeated LPS stimulations in our experimental model of tolerance. Interestingly, GM1 treatment also resulted in the down-regulation of a prototypical non-tolerizeable gene, *Fpr1*, in both Q7/Q7 and Q140/140 microglia. This suggests that rather than restoring gene silencing of tolerizeable genes in Q140/140 microglia, cell pre-incubation with the ganglioside might block stimulation by LPS altogether, as also shown in our previous studies in wild-type microglia (53).

The exact mechanisms underlying the anti-inflammatory effects of GM1 in normal and HD microglia awaits clarification. Our previous studies showed that GM1 administration decreases the activation of both NFkB and MAPK pathways required for pro-inflammatory cytokine expression and secretion (109-112), without significantly altering TLR4 levels (53). Furthermore, GM1 exerts its effects even after microglia incubation with LPS, suggesting that it must attenuate signaling downstream of TLR4 activation. We showed that both the ceramide tail of gangliosides and the specific composition of the glycan head group, including the presence of sialic acid, are required to mediate anti-inflammatory effects (53), suggesting that glycan-binding proteins, in particular sialic-acid binding proteins, might interact with gangliosides in a glycan-specific manner to mediate their signaling effects. Another potential mechanism could be through NFkB sequestration in lipid rafts, which represses NFkB signaling (113). The effects of GM1 pre-incubation (24h recovery period) in our experimental model of tolerance are particularly profound and might not necessarily occur with the same mechanism that dampens the expression of inflammatory molecules when GM1 is administered for a shorter time after cell stimulation with LPS. Gangliosides were shown to upregulate the expression of IRAK-3 in monocytes(72). IRAK-3 is a negative regulator of TLR4 signaling (114) that is involved in endotoxin tolerance (73, 74, 115) and the epigenetic suppression of tolerizeable genes (116). In our studies, Irak3 expression was not affected by GM1 in our tolerance model or in naïve microglia incubated solely with GM1, suggesting that a different, IRAK-3-independent mechanism underlies the effects of GM1 on microglia activation and reactivation.

In summary, our studies suggests that expression of mutant HTT in HD microglia does not result, *per se*, in a heightened microglia response to inflammatory stimuli, at least in a model of adult-onset HD, but causes a cell-autonomous impairment in the development of tolerance that might enable chronic inflammation in the brain and, in turn, contribute to disease progression. GM1 administration exerts potent anti-inflammatory effects that might be beneficial in HD patients to decrease neuroinflammation.

**Supplementary Fig. 1. Q7/7 and Q140/140 microglia express similar levels of TLR4 and TLR2 at the plasma membrane in naïve and stimulated conditions.** (A) Schematic representation and timeline of cell treatment with LPS. Representative histograms and relative flow cytometry quantification (% TLR4^+^-cells and median fluorescence intensity) of plasma membrane TLR4 in naïve microglia (I), after 12 h of exposure to LPS (100 ng/ml) (II), and after LPS removal and 24 h of recovery in serum-free medium (III). N ≥ 5. A two-sided unpaired *t*-test was used to compare TLR4 levels between genotypes. (B) Plasma membrane TLR2 was measured by flow cytometry in naïve microglia (I) and after 6 h of LTA (10 ug/ml) stimulation (II). Representative histograms and quantification of TLR2^+^-cells and TLR2 median fluorescence intensity are shown in the bar graphs. N23. Two-way ANOVA with Tukey’s multiple comparisons test. Bars are means ± STDEV.

**Supplementary Fig. 2. LPS treatment does not significantly affect the survival of Q7/7 and Q140/140 microglia.** Representative images of Q140/140 microglia stained with Hoechst (blue) and PI (yellow) after incubation in serum-free medium for 24 h (top panels), and Metaxpress software masks (bottom panels) used for the automated quantification of cell nuclei and propidium iodide (PI)-positive cells (dead cells) by high-content microscopy analysis of cell death. Scale bar = 150 μm. The graph shows the % of PI-positive cells in microglia cultures treated with or without LPS (100 ng/ml) for 24 and 48 h. N=4. Two-way ANOVA with Tukey’s post test.

**Supplementary Fig.3: Necrotic N2a cells carrying mutant HTT induce higher microglial expression of pro-inflammatory cytokines compared to necrotic cells carrying wild-type HTT.** Q7/7 and Q140/140 microglia were incubated with necrotic N2a25Q (25Q, wild-type HTT) or N2a97Q (97Q, mutant HTT) cells for 4 h (1:2 microglia to necrotic cells ratio). Graphs show the fold-change of pro-inflammatory cytokine gene expression compared to the expression induced by necrotic N2a25Q cells. mRNA levels of the indicated cytokines were normalized over the geometric mean of three housekeeping genes (Normalization Index) (N ≥ 4). Ratio paired *t*-test. \**p*<0.05.

**Supplementary Fig. 4. Comparable levels of cell death in Q7/7 and Q140/140 microglia after exposure to LPS and recovery.** LDH enzymatic activity released in the culture medium due to cell death was measured in microglia cultures incubated with or without LPS (100 ng/ml) for 12 h (**A**) and after 24 h recovery in serum-free medium. N=3. (**B)** Two-way ANOVA with Tukey’s multiple comparisons test. Bars are means ± STDEV*. *p<*0.05.

**Supplementary Fig. 5: GM1 decreases expression and production of pro-inflammatory cytokines in Q7/7 microglia.** Q7/7 microglia were activated with LPS (100 ng/ml) for 3 h, washed and treated with GM1 (50 µM) for 6 h. GM1 reduced the levels of **(A)** *Il-1b* and *Tnf* mRNA (N=5), and **(B)** TNF secreted in the medium (N ≥ 3). Gene expression was normalized over *Ppia*. Two-way ANOVA with Tukey’s multiple comparisons test was used. *\*p<0.05; **p<0.01; ***p<0.001, ****p<0.0001*.

**Supplementary Fig. 6: GM1 does not affect Irak-3 expression in naïve Q7/7 and Q14/140 microglia.** Naïve microglia were incubated with GM1 in serum-free medium for 8 h prior to RNA extraction and analysis of Irak-3 mRNA levels. Irak-3 expression was normalized over the geometric mean of three housekeeping genes. No statistically significant differences were detected among groups. N ≥ 3. Bars are means ± STDEV. Two-way ANOVA with Tukey’s multiple comparisons test.

## DECLARATIONS

### Ethical Approval and Consent to participate

All procedures on animals were approved by the University of Alberta Animal Care and Use Committee (AUP00000336) and were in accordance with the guidelines of the Canadian Council on Animal Care.

### Consent for publication

Not applicable.

### Availability of supporting data

All data generated or analysed during this study are included in this published article and its supplementary information files.

### Authors’ contributions

NS, SS and DG designed research and experiments; NS, DG and AZ performed experiments and analyzed data; SS supervised experiments and data analysis. NS and SS wrote the manuscript; all authors read and approved the final manuscript.

## LIST OF ABBREVIATIONS

FDR: Formyl peptide receptor 1
HD: Huntington’s Disease
IL-1β: Interleukin-1β
IL-6: Interleukin-6
IRAK-3: Interleukin 1 receptor associated kinase 3
LDH: Lactic dehydrogenase
LPS: Lipopolysaccharide
MAPK: Mitogen-activated protein kinase
NFκB: Nuclear factor NF-kappa-B
NO: Nitric Oxide
TNF: Tumor necrosis factor
TLR-4: Toll-like receptor 4
TLR-2: Toll-like receptor 2

## Acknowledgements

Schematics were generated using BioRender.com.

