## Supplementary figures and figure legends for "Naïve Huntington’s disease microglia mount a normal response to inflammatory stimuli but display impaired development of innate immune tolerance that can be counteracted by ganglioside GM1"

**Supplementary Fig. 6: GM1 does not affect *Irak-3* expression in naïve Q7/7 and Q14/140 microglia.** Naïve microglia were incubated with GM1 in serum-free medium for 8 h prior to RNA extraction and analysis of Irak-3 mRNA levels. Irak-3 expression was normalized over the geometric mean of three housekeeping genes. No statistically significant differences were detected among groups. N ≥ 3. Bars are means ± STDEV. Two-way ANOVA with Tukey’s multiple comparisons test.

### Supplementary Fig. 1

A

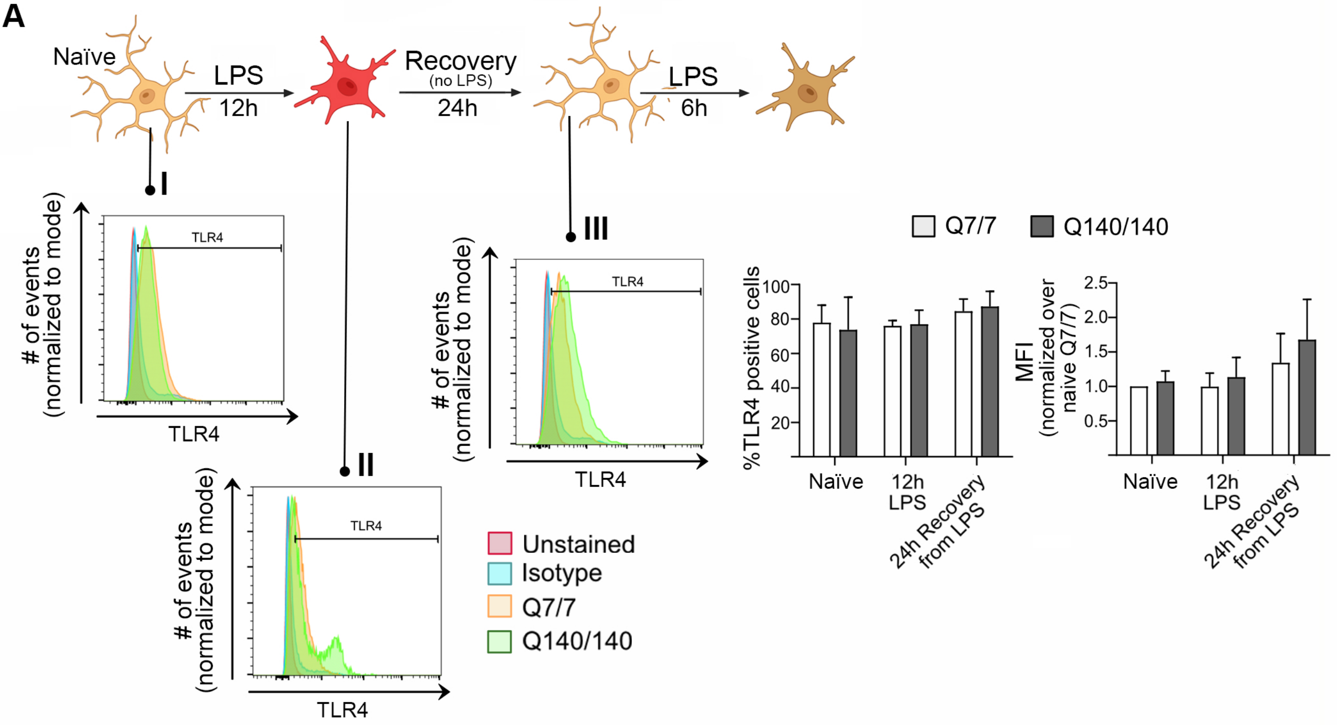


B

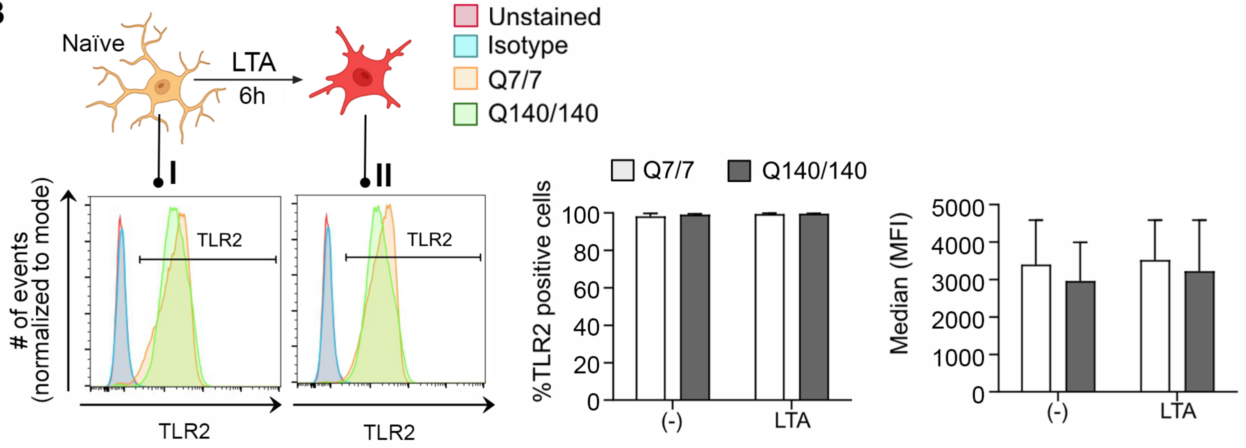

**Supplementary Fig. 2**

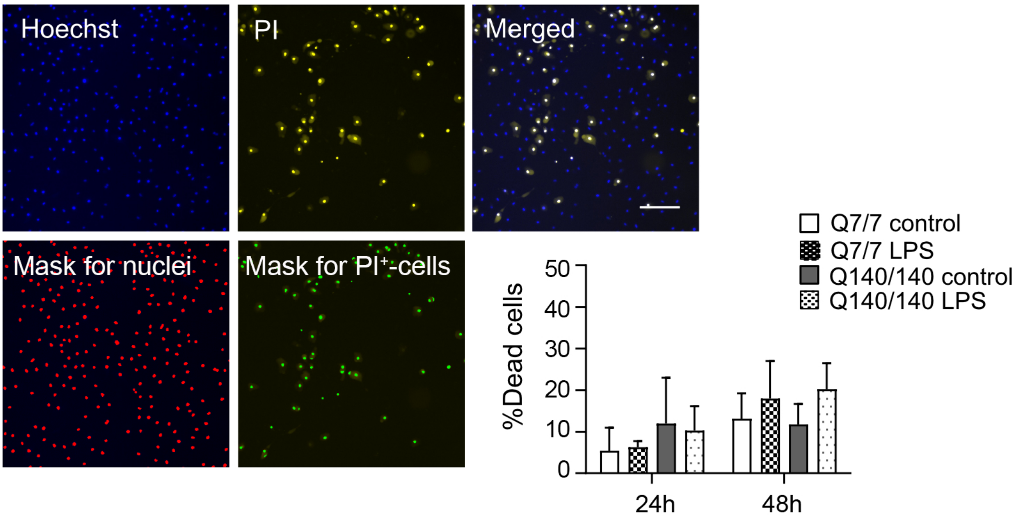

**Supplementary Fig. 3**

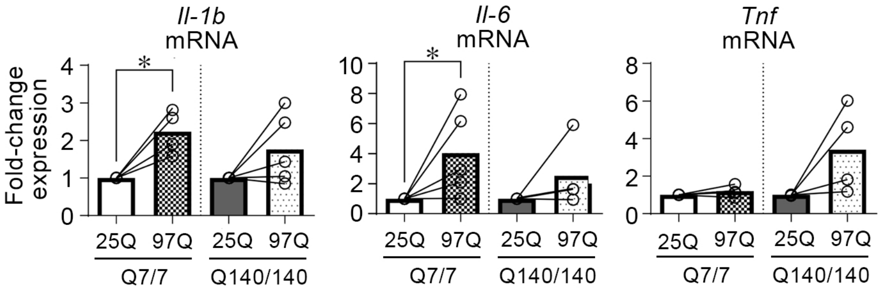

**Supplementary Fig. 4**

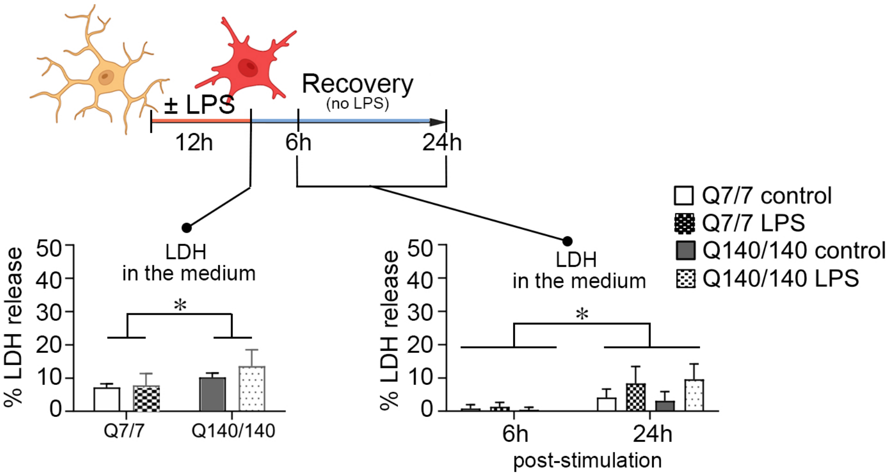

#### Supplementary Fig. 5

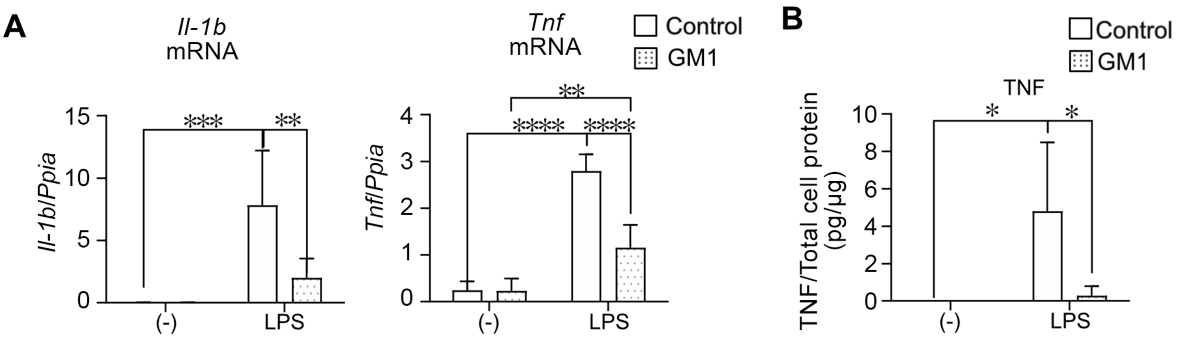

**Supplementary Fig. 6**

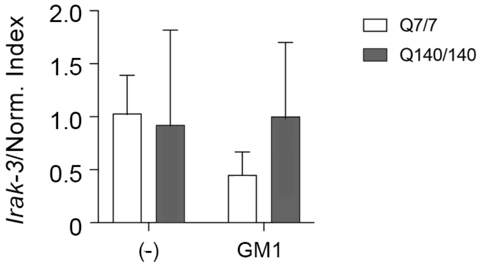
